# Amyotrophic lateral sclerosis associated mislocalisation of TDP-43 to the cytoplasm causes cortical hyperexcitability and reduced excitatory neurotransmission in the motor cortex

**DOI:** 10.1101/2020.06.11.147439

**Authors:** MS Dyer, KE Lewis, AK Walker, TC Dickson, A Woodhouse, CA Blizzard

**Affiliations:** Menzies Institute for Medical Research, College of Health and Medicine, University of Tasmania, Hobart, Tas, 7000, Australia; Neurodegeneration Pathobiology Laboratory, Queensland Brain Institute, University of Queensland, Brisbane, Queensland, 4072, Australia; Wicking Dementia Research and Education Centre, College of Health and Medicine, University of Tasmania, Hobart, Tas, 7000, Australia

**Keywords:** ALS, excitability, TDP-43, mislocalisation, cortex, mouse model, glutamate, hyperexcitability

## Abstract

Amyotrophic lateral sclerosis (ALS) is a chronic neurodegenerative disease pathologically characterised by mislocalisation of the RNA binding protein TAR-DNA binding protein 43 (TDP-43) from the nucleus to the cytoplasm. Changes to neuronal excitability and synapse dysfunction in the motor cortex are early pathological changes occurring in people with ALS and mouse models of disease. To investigate the effect of mislocalized TDP-43 on the function of motor cortex neurons we utilised mouse models that express either human wild-type (TDP-43^WT^) or nuclear localization sequence-deficient TDP-43 (TDP-43^ΔNLS^) on an inducible promoter that is restricted to the forebrain. Pathophysiology was investigated through immunohistochemistry and whole-cell patch-clamp electrophysiology. Thirty days expression TDP-43^ΔNLS^ in adult mice (60 days of age) does not cause any changes in the number of NeuN positive nor CTIP2 positive neurons in the motor cortex. However at this time-point the expression of TDP-43^ΔNLS^ drives intrinsic hyperexcitability in layer V excitatory neurons of the motor cortex. This hyperexcitability occurs concomitantly with a decrease in excitatory synaptic input to these cells. This pathophysiology is not present when TDP-43^WT^ expression is driven, demonstrating that the localisation of TDP-43 to the cytoplasm is crucial for the altered excitability phenotype. This study has important implications for the mechanisms of toxicity of one of the most notorious proteins linked to ALS, TDP-43. We provide the first evidence that TDP-43 mislocalization causes aberrant synaptic function and a hyperexcitability phenotype in the motor cortex, linking some of the earliest dysfunctions to arise in people with ALS to mislocalisation of TDP-43.

## Introduction

ALS is the most common adult-onset form of motor neuron disease, affecting the corticomotor system, which is comprised of upper motor neurons of the motor cortex and brain stem and lower motor neurons of the spinal cord. The symptoms are characterised by progressive muscle weakness, paralysis, and death from respiratory failure within 3-5 years of symptom onset. There are currently no effective treatments for the disease [1].

The hallmark pathological change seen in post-mortem tissue from both sporadic (~90% of cases) and familial (~10%) ALS patients is the cytoplasmic accumulation and aggregation of the RNA/DNA-binding protein, transactive response DNA-binding protein of 43 kDa (TDP-43) [2, 3]. The primary role of TDP-43 is to regulate the degradation, localization and splicing of RNA and under normal conditions TDP-43 is tightly autoregulated [4]. Whilst predominately nuclear, a small amount of TDP-43 is present in the cytoplasm, and plays important roles in the maintenance of excitatory synaptic structure and function [5]. An enduring research question remains regarding the contribution of TDP-43 to ALS; what is the functional consequence of mislocalised TDP-43 to the cytoplasm?

One of the earliest detectable physiological changes in ALS are motor cortex excitability disturbances. This has been observed clinically with the use of transcranical magnetic stimulation (TMS), a non-invasive depolarisation technique. TMS studies of the ALS motor cortex have observed a reduced resting motor threshold (measurement of intrinsic excitability of corticospinal motor neurons) at early disease stages [6, 7]. Furthermore, increased intracortical facilitation, a measurement of glutamatergic function, is also increased in ALS [8]. Cortical hyperexcitability of the motor cortex has been detected in sporadic ALS patients prior to spinal motor neuron dysfunction and in carriers of a SOD1 mutation conferring familial ALS before symptom onset [9-11]. These alterations in excitability are present in many mouse models of familial ALS, including those with familial TDP-43 transgenes, long before the onset of disease-like symptoms [12-16]. In the SOD1 mouse model of ALS, cortical neuron hyperexcitability is observed in neonates, before normalising and eventually becoming hypoexcitable at a late disease stage [14] Excitability changes are a consistent feature in cases, as well as mouse and other animal models of familial ALS, yet the initial cause of cortical hyperexcitability in sporadic ALS remains unknown.

This study aimed to interrogate if TDP-43 mislocalization alters the physiology of neurons which are vulnerable in ALS. In this study we used a mouse model where human nuclear localization sequence-deficient TDP-43 (TDP-43^ΔNLS^) expression was restricted to the forebrain with the calcium/calmodulin dependant protein kinase II alpha (CAMKIIα) promoter and was temporally induced with the tetracycline transactivator system [17]. This model expresses TDP-43 that is mislocalised but does not contain familial mutations, modelling the pathology that occurs in sporadic ALS. Furthermore, a secondary model which expresses TDP-43 (TPD-43^WT^) with a functioning nuclear localisation sequence on the same promoter and tetracycline transactivator system was utilised to determine if increased levels TDP-43 alone, rather than mislocalisation, contributed to altered excitability. We present the first evidence that TDP-43 mislocalisation in the motor cortex causes hyperexcitability of upper motor neurons and decreases cortical excitatory neurotransmission. These data demonstrate that TDP-43 mislocalisation causes upper motor neuron hyperexcitability, linking the hallmark cytoplasmic TDP-43 pathology with a major early physiological change that occurs in ALS.

## Methods

### Mouse cohort

All experiments were approved by the Animal Ethics Committee of the University of Tasmania (#A16593) in accordance with the Australian Code of Practice for the Care and Use of Animals for Scientific Purposes (2013), was exploratory and was not pre-registered. Mice had ad libitum access to food and water and were housed in groups of up to five but were not singly housed. CamkIIα-tTA mice were bred to a previously described tetO-hTDP-43-ΔNLS line 4 mice (TDP-43^ΔNLS^) and tetO-hTDP-43-WT line 12 mice (TDP-43^WT^) [17]. All lines were fully back-crossed to a pure C57BL/6J background prior to commencing experiments. Monogenic CamkIIα-tTa were used as controls. Breeding mice were provided with chow containing 200mg/kg doxycycline (dox) at least 1 week before mating and their offspring remained on this diet until postnatal day 30 (P30). At P30 mice were switched to standard chow, inducing expression of the transgene and aged to P58-P63 (28-33 days on standard chow), a timepoint which has previously been reported to have significant motor and cognitive behavioural changes [18]. Male mice were used for electrophysiological experiments. Sample sizes for experiments were empirically determined.

### Immunohistochemistry for cell counts

For quantification of TDP-43 misloclisation in cortical excitatory neurons, five 40 μM slices containing motor cortex were selected from five TDP-43^ΔNLS^ mice each. These free-floating sections were blocked with 1% normal goat serum for 1 hour in 0.01M phosphate buffered saline (PBS) before incubation with with a mouse α-NeuN (Fox-3) antibody (1:1000; Millipore, Cat#: MAB377 (2018)) and a rabbit α-TDP-43 antibody (1:1000; Proteintech, Cat#: 10782-2-AP (2018)) overnight in PBS with 0.3% triton-X. Secondaries (Alexa Fluor anti-rabbit 568 and anti-mouse 488, 1:1000, Molecular Probes) were applied for 90 minutes. All incubations were performed at room temperature. Sections were mounted onto slides and coverslipped with PermaFluor aqueous mounting media (Thermo-Fisher Scientific).

For quantification of COUP-TF-interacting protein-2 (CTIP2)-positive projection neurons, slices containing primary motor cortex from bregma 1.78 to 0.14 as from Paxinos and Franklin mouse brain atlas [19] were arranged in rostral-caudal order and every 5^th^ 40μM slice chosen for counting (eight slices per animal). Five WT, five TDP-43^WT^ and five TDP-43^ΔNLS^ mice (one mouse from this group was deemed uncountable due to vessel staining) were used for experiments. Endogenous peroxidase activity was quenched by incubation with a mixture of 3% H_2_O_2_ and 10% methanol in PBS. Sections were then incubated with a rat α-CTIP2 (1:500; Abcam, Cat#: ab18465 (2019)) overnight at room temperature. After rinsing, sections were incubated with a biotinylated secondary antibody solution (1:200, goat α-rat, Vector Laboratories) for 1 hour. Following treatment with avidin-biotin-peroxidase kit (Vector Laboratories), colour reaction was developed by using a 3,3’-diaminobenzidine kit (Vector Laboratories). Sections were mounted on slides, dried and coverslipped with DPX mounting media (Sigma-Aldrich).

### Microscopy

Images were acquired on an Andor spinning disk microscope and a 20x objective with NIS-elements software (Nikon). Imaging cohorts for counting analysis were imaged with identical microscope settings and all images were processed and prepared for presentation using ImageJ (NIH) and the Adobe Photoshop and Illustrator programs.

### Immunohistochemical Quantification

Quantification of NeuN and TDP-43 labelling was performed in Image J (NIH) and manually counted to a standardised area of 31250 μm^2^. Regions of interest (motor cortex vs somatosensory cortex, layer V vs layer II/II) were identified using the Paxinos and Franklin mouse brain atlas [19].

Quantification of CTIP2 labelling was performed first by selecting the motor cortex from the corpus callosum to the layer IV/V boundary layer from every fifth section [19] and using Image J to form a standardised grid of 40μM^2^. Every third box was counted to give an estimation of one-third of the number of cells in each slice, which was then corrected at the end for total number of CTIP2-positive cells in each slice of motor cortex. The experimenter was blinded to genotype at the image quantification stage.

### Slice Preparations

Male mice were anesthetized with an intraperitoneal injection of 150μL of 60mg/mL sodium pentobarbitone for ethical anesthetisation and transcardially perfused; brains were sliced (400μm coronal sections, Leica VT1200s vibratome) in ice cold, carbogenated (95% O_2_, 5% CO_2_) perfusion solution containing (in mM): 92 choline chloride, 2.5 KCl, 1.2 NaH_2_PO_4_, 30 NaHCO_3_, 20 HEPES, 25 Glucose, 0.5 CaCl_2_, 10 MgSO_4_.7H_2O_, 5 sodium ascorbate, 2 Thiourea, 3 sodium pyruvate 5 N-Acetyl-L-Cysteine [20]. Slices were then recovered for 30 minutes in a holding solution at 31°C containing (in mM): 92NaCl, 2.5KCl, 1.2NaH_2_PO_4_, 30 NaHCO_3_, 20 HEPES, 25 Glucose, 2 CaCl, 2MgSO_4_.7H_2_O 5 sodium ascorbate, 2 Thiourea, 3 sodium pyruvate 5 N-Acetyl-L-Cysteine [20]. Following recovery, slices were incubated at room temperature in carbogenated aCSF (125 NaCl, 25 NaHCO_3_, 2.5 KCl, 1.25 NaH_2_PO_4_, 2 CaCl_2_, 2 MgCl_2_ and 25 glucose, all from Sigma Aldrich).

### Electrophysiology

Patch pipettes were pulled from borosilicate glass capillaries (Harvard Apparatus) on a Sutter P1000 pipette puller (Sutter Instruments, USA) giving a tip resistance of 3-5MΩ and filled with intracellular solution with added Neurobiotin (~0.1%; Vector Labs) to allow identification of Layer V pyramidal neurons using post-hoc streptavidin labelling. All patched cells in transgenic mice were identified to express human TDP-43 (Fig 3A). Candidate layer V pyramidal neurons were identified through a 40X water immersion objective in a chamber continuously perfused (>4mL/min) with aCSF at 30°C (±1 °C). Whole-cell patch-clamp recordings of layer V pyramidal neurons in the M1 region of motor cortex with an uncompensated series resistance of <20 MΩ were included in the study. A liquid junction potential of +13.2mV was compensated for online. Recordings were acquired at 20kHz and were filtered with a low pass filter at 3kHz. Current and voltage clamp protocols were made and recorded with Patchmaster software (Heka Elektronik) and a Heka EPC10 USB amplifier (Heka Elektronik).

**Fig 3.**
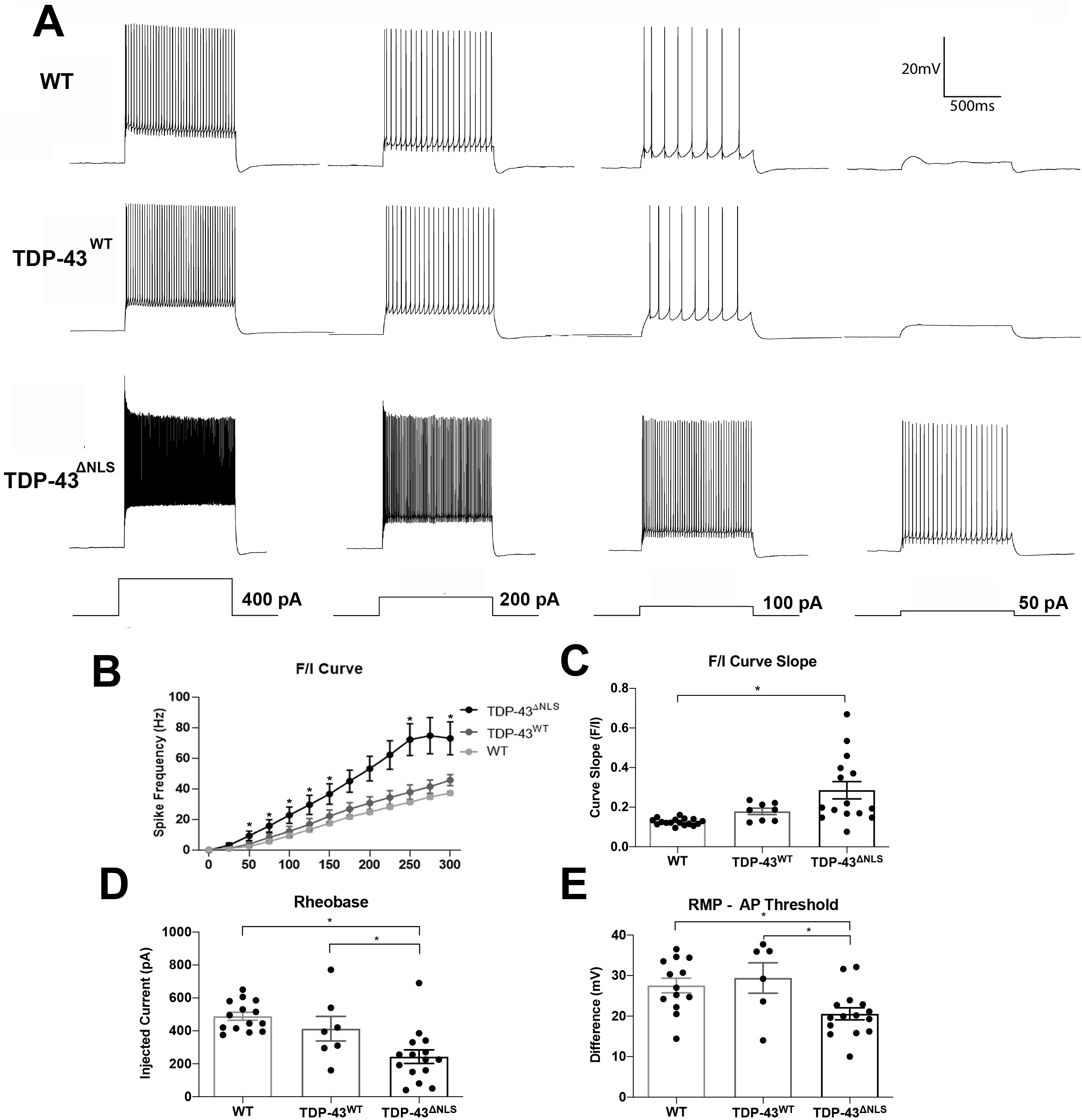
TDP-43^ΔNLS^ layer V excitatory neurons in the motor cortex are hyperexcitable at 30 days TDP-43 expression. A. Representative traces of WT, TDP-43^WT^ and TDP-43^ΔNLS^ neurons in response to 1 second stepped current injections of 400pA, 200pA, 100pA and 50pA respectively. B. Firing frequency of TDP-43^ΔNLS^ neurons was increased compared to WT controls in response to 400pA and 200pA current injections but no difference was observed at 100pA and 50pA current steps. C. There was an increase in the slope of the frequency/injected current (F/I) curves in TDP-43^ΔNLS^ neurons compared to WT neurons. D. TDP-43^ΔNLS^ neurons had a lower rheobase compared to both WT and TDP-43^WT^ cells. E. The difference between the resting membrane potential and the action potential threshold was smaller in TDP-43^ΔNLS^ neurons compared with WT and TDP-43^WT^ neurons. Each data point represents one cell.

Passive membrane properties and intrinsic excitability recordings were performed with an intracellular solution containing (in mM) 120 K-gluconate, 20 KCl, 10 HEPES, 10 Ethylene glycol-bis(2-aminoethylether)-N,N,N’,N’-tetraacetic acid (EGTA), 2 MgCl2, 2 Na2 and 0.3 NaGTP, pH 7.4, 280-290mOsm (Sigma Aldrich), and with 100 μM picrotoxin (Abcam) and 2mM kynurenic acid (Sigma Aldrich) in the bath to isolate the cell from synaptic activity. Passive membrane properties (τ_m_, R_in_, C_m_) were measured with 0.5s negative current pulses inducing small hyperpolarisations (<10mV). 1s incremental depolarising current steps from 0 to 400pA (25pA steps) were injected to produce action potential (AP) trains and the frequency vs current (F/I) curve was generated by using average frequency for each group. Rheobase was determined by the injection of depolarising current for 5ms with incremental 5pA steps. Action potential (AP) properties (threshold, half width, amplitude, rise/decay time and slope and afterhyperpolarisation) were determined from single spikes from the rheobase protocol. AP threshold was defined as the membrane voltage value reached when the voltage to first derivative rises to 5% of its peak amplitude. AP rise and decay slopes were measured between 10 and 90% of AP amplitude. Analysis of passive membrane properties, I/F curves, spike frequency and rheobase was performed using PatchMaster v2×90.4 (HEKA Elektronik), spike threshold in Spike 2 v7.10 (Cambridge Electronic Design) and action potential properties in pCLAMP (Molecular Devices). A total of 20 mice were used for intrinsic excitability experiments.

Excitatory inputs were recorded in aCSF in the presence of picrotoxin to eliminate GABA-ergic currents with an intracellular solution consisting of (in mM) 120 Cs-methanesulfunate, 20 CsCl, 10 HEPES, 10 EGTA, 2MgCl_2_, 2Na2ATP and 0.3 NaGTP, pH 7.4, 280-290mOsm (Sigma Aldrich)., Voltage-clamp recordings were performed at −70mV (no liquid junction potential correction) for 2 minutes using glass pipettes with a tip resistance of 3-4 MΩ. mEPSCs were recorded with the addition of tetrodotoxin (Abcam) to the bath. The baseline was manually adjusted offline if not completely level and inputs that were more negative than 10mV were counted using clampfit 10.7. All recordings had a series resistance of <15 MΩ and noise of <10pA, only recordings with a stable series resistance over the 2-minute recording were used for analysis. No blinding was performed for electrophysiological experiments.

### Immunohistochemistry of Neurobiotin cells

After filling with Neurobiotin, the slice was fixed in 4% paraformaldehyde in 0.1M phosphate buffer pH 7.4 overnight. The sections were incubated for 1 hour in a blocking solution containing 0.2% bovine serum albumin, 1% normal goat serum and 0.5% triton-X100 in 0.1M phosphate buffered saline. Alexa Fluor 568 streptavidin (1:1000, Molecular Probes, Cat#: S11226), diluted in 0.2% bovine serum albumin, 0.2% normal goat serum and 0.5% triton-X100 was then applied overnight before washing in 0.1M phosphate buffered saline and mounted for imaging.

### Statistical Analysis

All data were analysed in GraphPad Prism version 8. One-way analysis of variance (ANOVA) with post-hoc Bonferroni correction was employed to compare between groups. A p-value of <0.05 was considered statistically significant. No tests for outliers were conducted on data.

## Results

### Thirty days expression of TDP-43^ΔNLS^ in neurons of the forebrain does not cause overt cell death of projection neurons in the motor cortex

Here we utilise two mouse models with TDP-43 alterations, the TDP-43^ΔNLS^ model has wildtype TDP-43 expressed in forebrain neurons under control of the CamkIIα promoter, but with a dysfunctional nuclear localisation sequence, causing cytoplasmic localisation of TDP-43 [17]. The TDP-43^WT^ mouse model expresses wild-type TDP-43 on the same promoter with a functioning nuclear localisation sequence, and the TDP-43 in these cells is mostly nuclear [2]. The mouse models used in these experiments were first fully back-crossed (10 generations) to the C57Bl6 strain; which shows minimal physiological side-effects of tetracycline-transactivator protein expression [21]. Immunohistochemistry with antibodies directed at human TDP-43, and NeuN in the cortex of TDP-43^ΔNLS^ mice at 30 days expression confirmed that human TDP-43 was expressed in the cytoplasm of excitatory neurons of the neocortex (Fig 2A, B), with ~80% of NeuN positive cells in the motor cortex and somatosensory cortex being positive for human TDP-43 (Fig 2C). Of the human TDP-43 positive cells, 100% were positive for NeuN, indicating that expression was restricted to neurons (Fig 2D). Thirty days of TDP-43^ΔNLS^ expression did not result in a significant loss of CTIP2 positive neurons, a marker for projection neurons in layers V and VI including motor neurons [22, 23](Fig 2E). At this timepoint, previous studies have shown both motor and cognitive dysfunction in the TDP-43^ΔNLS^ mouse model [17, 18], indicating dysfunction of the motor cortex without overt cell death.

**Fig 1.**
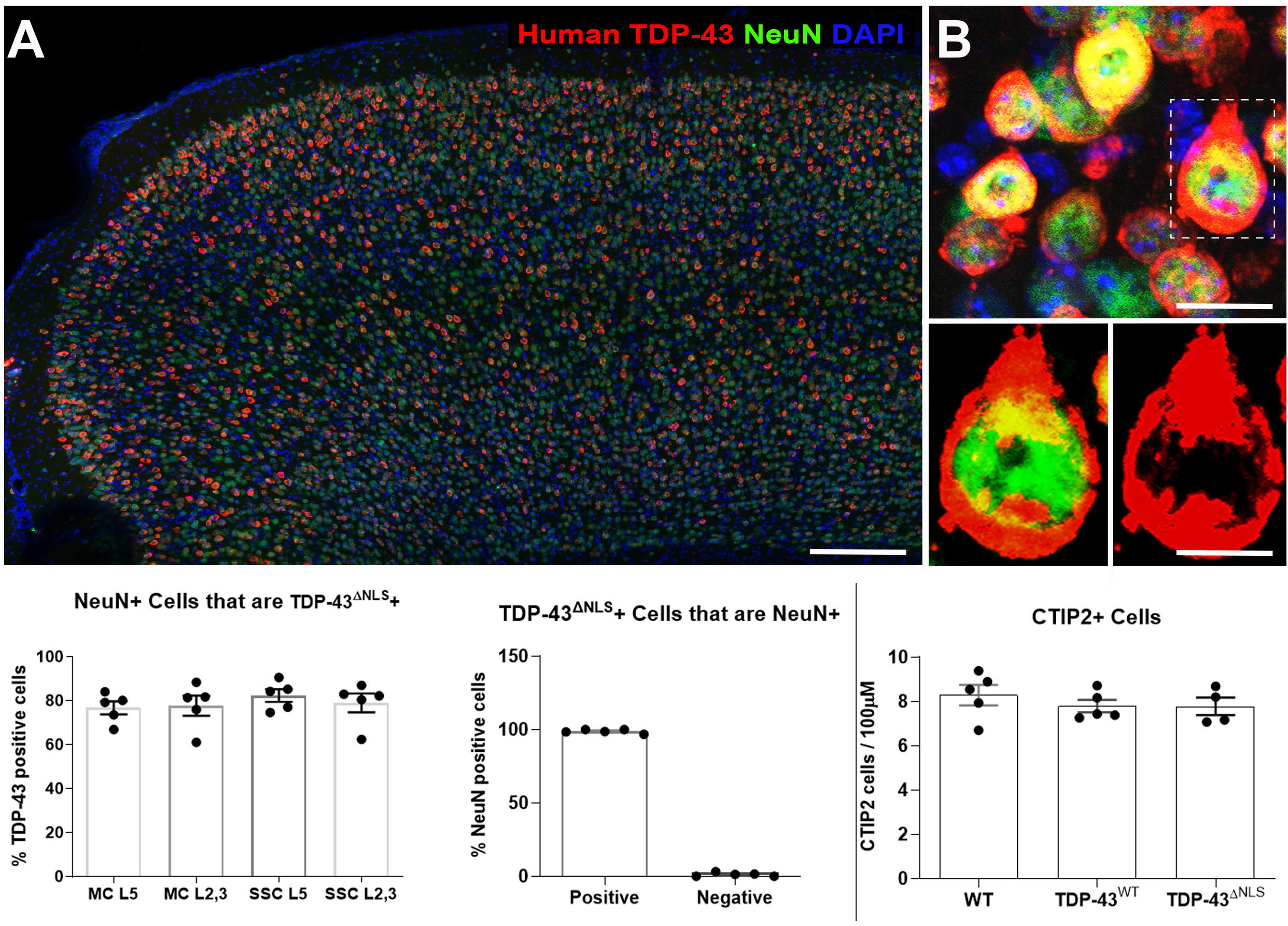
TDP-43^ΔNLS^ is expressed in neurons throughout the cortex, but does not cause loss of CTIP2-positive projection neurons from the motor cortex. A. Immunohistochemistry of a TDP-43^ΔNLS^ mouse cortex labelled with hTDP-43 (red), NeuN (green) and DAPI (blue) shows that hTDP-43 is expressed throughout the forebrain (scale bar 50μm). B. hTDP-43 (red) is mislocalized out of the nucleus to the cytoplasm in a subset of NeuN (green) positive cells (scale bar 50μm). Inset shows TDP-43 surrounding the nucleus of a NeuN labelled cell (scale bar 20μm). C. The proportion of NeuN cells positive for hTDP-43 in the motor and somatosensory cortices in cortical layer 2/3 and 5 is ~80%. D. Almost every cell positive for TDP-43 was also positive for NeuN. E. There was no significant difference in the number of CTIP2-positive neurons between genotypes. Each data point represents the average between slices of one animal.

**Fig 2.**
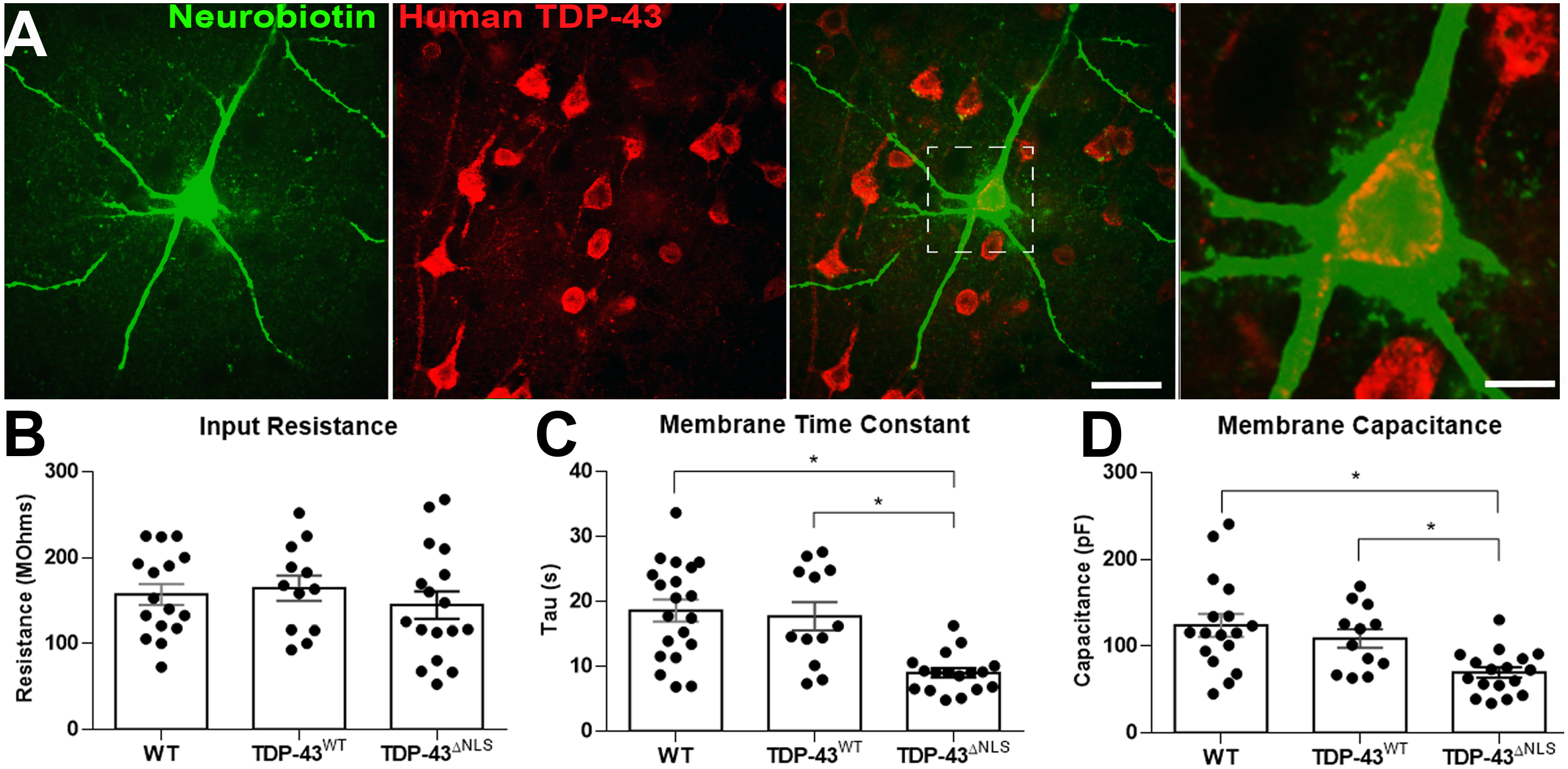
TDP-43^ΔNLS^ layer V pyramidal neurons are altered in their electrophysiological membrane properties. A. Layer V pyramidal neurons of the motor cortex were selected for patch clamp recording and filled with Neurobiotin (green). Post hoc double labelling with an antibody to hTDP-43 (red) confirmed that the neurons selected for electrophysiological analysis were positive for TDP-43 in the TDP-43^ΔNLS^ cortex (scale bar 50μm). B. There were no changes to input resistance. C. There was a significantly faster membrane time constant in TDP-43^ΔNLS^ neurons compared to WT and TDP-43^WT^ (p<0.05). D. TDP-43^ΔNLS^ motor cortex neurons had a decrease in membrane capacitance compared to WT and TDP-43^WT^ neurons (p<0.05). Each data point represents one cell.

### Layer V pyramidal neurons in the motor cortex exhibit mislocalised TDP-43 and altered passive membrane properties

We next investigated if mislocalized TDP-43 influences the fundamental membrane properties of neurons. Layer V pyramidal neurons of the motor cortex are particularly vulnerable to cell death in ALS and have early TDP-43 accumulation in disease [24-26]. The passive membrane properties of layer V pyramidal neurons of the M1 motor cortex were measured with whole-cell patch-clamp electrophysiology. TDP-43^ΔNLS^ (Fig 3A), and TDP-43^WT^ layer V pyramidal neurons exhibited no change to input resistance (Fig 3B), but TDP-43^ΔNLS^ cells showed a significantly decreased membrane time constant (Fig 3C) and membrane capacitance (Fig 3D) compared to both TDP-43^WT^ mice and WT mice.

### Layer V pyramidal neurons in the motor cortex with cytoplasmic TDP-43 mislocalisation are hyperexcitable

To determine the impact of mislocalisation of TDP-43 in upper motor neurons we investigated the intrinsic excitability of M1 motor cortex layer V pyramidal neurons. When the responses to depolarizing current injections (Fig43A) were analysed, TDP-43^ΔNLS^ layer V pyramidal neurons exhibited increased firing rates (Fig 4B), corresponding to an increase in the frequency/injected current (F/I) curve slope (Fig 4C) compared to TDP-43^WT^ and WT neurons. There was no significant change to the intrinsic excitability of TDP-43^WT^ neurons. TDP-43^ΔNLS^ neurons showed a lower rheobase compared to both WT and TDP-43^WT^ neurons (Fig 4D), indicating that they require less excitatory stimulation to fire action potentials. The origin of the altered rheobase in TDP-43^ΔNLS^ layer V pyramidal neurons can be at least partly explained by a reduction in the difference between the resting membrane potential (RMP) and the action potential threshold (Fig 4E). There were no changes to rheobase or RMP – action potential threshold in the TDP-43 ^WT^ neurons (fig 4D, E). Collectively these data indicate that the mislocalisation of TDP-43 to the cytoplasm causes the hyperexcitability of layer V pyramidal neurons.

**Fig 4.**
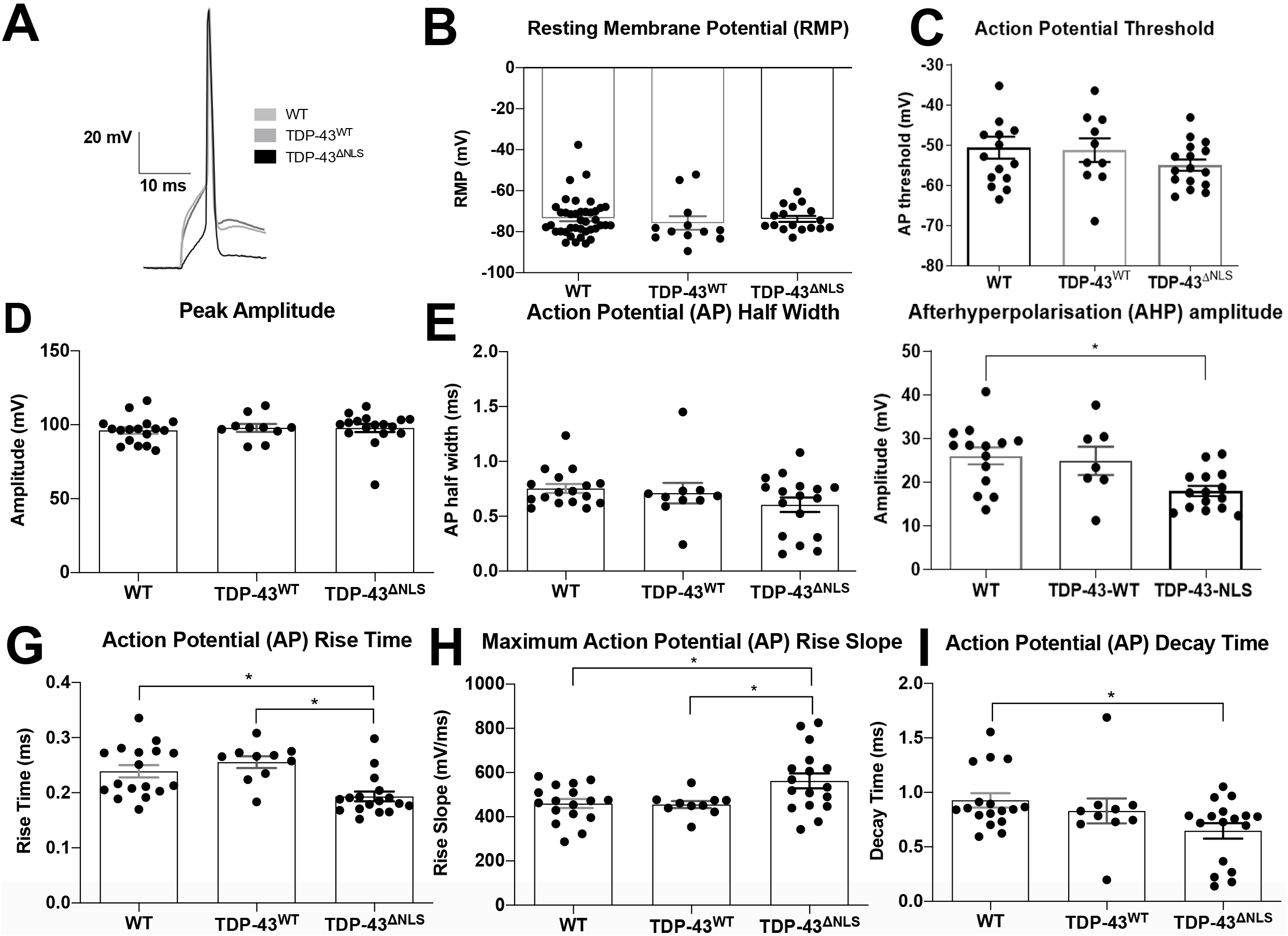
TDP-43^ΔNLS^ layer V excitatory neurons in the motor cortex have alterations to action potential waveforms. A. Representative action potentials from WT (light grey) TDP-43^WT^ (grey) and TDP-43^ΔNLS^ (black) neurons at rheobase. B. There was no change to the resting membrane potential between the three groups. C. There was no significant change to the action potential threshold. D. There was no difference in AP peak amplitude in the TDP-43^ΔNLS^ neurons in comparison to both WT and TDP-43^WT^ neurons. E. There was no difference in AP half-width of neurons between genotypes. F. TDP-43^ΔNLS^ neurons have a decreased AHP compared to WT layer V pyramidal neurons (p<0.05). G. TDP-43^ΔNLS^ neurons have a faster AP rise time than both WT and TDP-43^WT^ cells and have a greater maximum slope of the rising portion of the action potential (p<0.05, H,I). I. The AP decay time was also decreased in TDP-43^ΔNLS^ neurons, in comparison to WT (p<0.05). Each data point represents one cell.

### TDP-43^ΔNLS^ expressing neurons in layer V of the motor cortex demonstrate a smaller afterhyperpolarisation (AHP) and altered action potential kinetics

To see if hyperexcitability might be caused by ion channel alterations that could alter action potential waveform, action potential characteristics (Fig 5A) were analysed. There was no difference in the RMP (Fig 5B) or the action potential threshold (Fig 5C), peak amplitude (Fig 5D), half width (Fig 5E) and maximal decay slope (data not shown) between groups. However, the AHP amplitude was significantly decreased in the TDP-43^ΔNLS^ neurons (Fig 5F), and there was a small but significant decrease in the action potential rise time (Fig 5G) and increase in the maximal rise slope (Fig 5H) in comparison to both WT and TDP-43^WT^ neurons. Decay time was also significantly decreased in TDP-43^ΔNLS^ vs WT, however this difference was not significant between TDP-43^ΔNLS^ and TDP-43^WT^ neurons (Fig 5I). Collectively this indicates that the action potential waveform is subtly altered in TDP-43^ΔNLS^ layer V pyramidal neurons while a decrease in the AHP amplitude may mean that the time required to fire second and subsequent action potentials is decreased in the presence of mislocalised TDP-43, contributing to the observed hyperexcitability.

**Fig 5.**
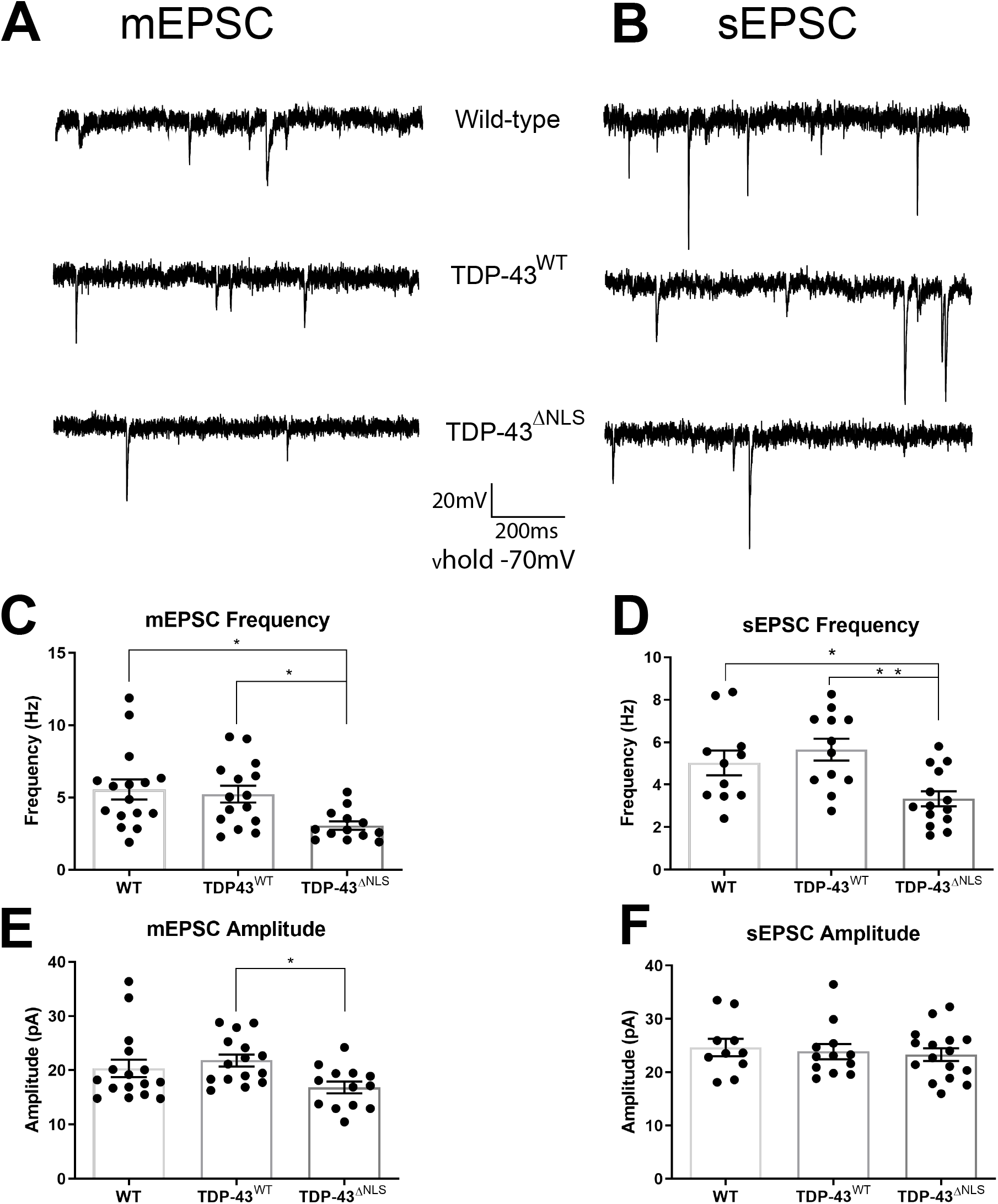
Motor cortex TDP-43^ΔNLS^ layer V neurons receive reduced excitatory input. A) Example traces of mini-excitatory post-synaptic currents (mEPSCs, no action potentials) from WT, TDP-43^WT^ and TDP-43^ΔNLS^ neurons. B) Example traces of spontaneous excitatory post-synaptic currents (sEPSCs, with action potentials). C) Frequency of mEPSCs is decreased in TDP-43^ΔNLS^ neurons (p<0.05). D) The frequency of TDP-43^ΔNLS^ sEPSCs are decreased compared to WT and TDP-43^WT^, E) The amplitude of mEPSCs are smaller in TDP-43^ΔNLS^ neurons compared to TDP-43^WT^, but not to WT neurons. E) There is no difference to the average sEPSC amplitude. Each data point represents one cell.

### Layer V pyramidal neurons expressing cytoplasmic TDP-43 have reduced excitatory synaptic inputs

Emerging evidence shows that synaptic dysfunction of cortical neurons is associated with ALS. In the TDP-43^ΔNLS^ mouse model, mislocalised TDP-43 is expressed in excitatory forebrain neurons and is uniformly expressed throughout both the layers and different areas of the cerebral cortex (Fig 2A, C). We next investigated if excitatory inputs were altered by TDP-43^ΔNLS^ expression. The frequency of mini excitatory post-synaptic currents (mEPSCs) to TDP-43^ΔNLS^ layer V pyramidal neurons was lower compared to both WT and TDP-43^WT^ cells (Fig 6C, p<0.05). Similarly, TDP-43^ΔNLS^ layer V pyramidal neurons also showed a decrease in the frequency of spontaneous excitatory post-synaptic currents (sEPSCs), with the presence of action potentials in the slice (Fig 6D, p<0.05). This suggests that there is either a reduced number of excitatory synapses on to layer V pyramidal neurons or a reduction in the release probability from the presynaptic compartment in TDP-43^ΔNLS^ neurons. There was a significant decrease in the amplitude of mEPSC compared to TDP-43^WT^, but not to WT neurons (Fig 6E), but no change to the amplitude of sEPSC (Fig 6F). Together these data show that TDP-43 mislocalisation in the forebrain causes a loss of excitatory neurotransmission to layer V pyramidal neurons in the primary motor cortex.

## Discussion

Alterations to neuronal excitation have long been associated with ALS, with increased concentrations of the excitatory neurotransmitter glutamate reported in the cerebrospinal fluid of ALS patients 30 years ago [27]. However, the underlying cause of neuronal hyperexcitability has still not been determined. Understanding how ALS progresses through the very first stages of symptoms, when patients are likely to present to a clinic, to rapidly become a debilitating disease is critical for the production of future therapeutics. Here, we show for the first time that TDP-43 pathologically located to the cytoplasm, as occurs in sporadic ALS [3], causes cortical neuronal hyperexcitability, a defining physiological feature of the disease [28, 29]. Our understanding of the pathogenesis of ALS has improved with the discovery of the significant contribution of the cortex to disease. Cortical hyperexcitability precedes neurodegeneration and the eventual neuro-motor neurotransmission failure that occurs in ALS [9, 10]. This study is the first to show that cortical neurotransmission failures before cell loss in ALS can be driven by the pathological mislocalisaiton of TDP-43 to the cytoplasm.

The electrophysiological findings presented here are consistent with human studies of ALS. We observe hyperexcitable layer V excitatory neurons in the motor cortex, as is seen in clinical TMS data [8-10]. While synaptic inputs cannot be measured non-invasively in humans, a marked decrease in dendritic spines and dendritic retraction has been observed in the human primary motor cortex of sporadic ALS cases, indicating a loss of synaptic input [30]. The similarities between our studies and observed human pathology could be due to the disease model used here having pathology that is more representative of sporadic ALS and thus a better model of the majority of ALS cases. Similar mechanisms may occur in familial ALS cases that harbour mutations to TDP-43; for example, sEPSC frequency but not amplitude is reduced in the TDP-43 A315T motor cortex in mice [15].

We show that TDP-43 mislocalisation drives hyperexcitability in layer V excitatory neurons in the motor cortex of TDP-43^ΔNLS^ mice in comparison to TDP-43^WT^ and their wild type littermate controls. This increased excitability was characterised by a decrease in rheobase, which could be accounted for by a reduced difference between the resting membrane potential (RMP) and action potential threshold. This could be caused by alterations to the conductances of a number of ion channels including voltage activated sodium and outward potassium currents. Our studies revealed alterations in the action potential rise time and rise slope in the TDP-43^ΔNLS^ neurons in comparison to wild type littermate controls, indicative of changes to voltage-gated sodium channels. Indeed, changes to voltage activated sodium channels have been previously observed in familial models of ALS [31, 32], suggesting this may be a conserved mechanism between sporadic and familial models. Furthermore, the fast and slow components of the AHP were reduced in TDP-43^ΔNLS^ cells suggesting alterations in voltage- and calcium-dependent potassium channels [33].

The first effective treatment for ALS, Riluzole, is proposed to reduce hyperexcitability by inhibiting the persistent sodium current, thus increasing the action potential threshold and returning hyperexcitable neurons towards a normal level of activity [34, 35]. Our data provide a potential mechanism for why Riluzole is effective in treating early sporadic disease - by protecting against the TDP-43 mediated alterations in voltage-gated sodium channels. Interestingly, altered intrinsic excitability is a complex and evolving phenomenon in ALS. A switch from hyper- to hypo-excitability occurs in the SOD1 G93A mouse model, potentially involving persistent sodium current alterations [14]. Hypoexcitability later in ALS progression may also contribute to the reduced effectiveness of Riluzole at later disease stages. How mislocalised TDP-43 affects neuronal excitability at different stages of the disease time course is an interesting question that warrants further consideration.

We revealed that intrinsic hyperexcitability occurs concurrently with decreased excitatory synaptic inputs to layer V pyramidal neurons. [12, 36]. The main excitatory pathway of excitatory input to motor cortex layer V pyramidal neurons is from the layer 2/3 population of the primary motor cortex [37]. Given that a large proportion of layer 2/3 neurons express mislocalised TDP-43 in this mouse model, and therefore might also be expected to exhibit hyperexcitability, it could be considered surprising that there are reduced sEPSCs to layer V pyramidal neurons. The reduction in mEPSC frequency observed in the current study suggest that there is either synapse loss or altered presynaptic release probability. If layer 2/3 neurons are indeed hyperexcitable due to expressing mislocalised TDP-43, synapse loss on layer V pyramidal neurons would explain the observed decrease in mEPSCs and sEPSCs. Though it may appear unexpected, these results are in line with electrophysiological data from the TDP-43 A315T familial ALS mouse model, which presents with increased intrinsic hyperexcitability of layer V pyramidal neurons from 3 weeks to 15 weeks of age, decreased frequency of mEPSC and reduced spine density (a proxy for measuring excitatory synaptic input) at ~9 weeks [15, 16]. It remains to be elucidated if concomitant increased intrinsic excitability and reduced excitatory synaptic inputs are specific to models of TDP-43 or shared between all ALS models. Spine loss has been observed in motor cortex layer V pyramidal neurons in symptomatic SOD1 G93A mice [38], though occurring concurrently with spine loss is an increase in synaptic excitation, probably due to the simultaneous hyperexcitability of other excitatory cortical neurons.

Another possibility for the mechanism of action of mislocalised TDP-43 is that it is causing either intrinsic hyperexcitability, synapse loss or reduced synaptic excitability and the neuron is maintaining a set point of activity through homeostatic plasticity [39-41]. Determining if one property is pathogenic and one is homeostatic however, remains a difficult problem for motor systems *in vivo*. At the same time, mislocalised TDP-43 influences the expression of thousands of genes, so it is also plausible that mislocalised TDP-43 could influence both excitatory synapses and intrinsic excitability independently [42, 43]. To add to the complexity of TDP-43 induced altered excitability in ALS, a recent study elegantly showed that driving hyperexcitability pharmacologically can cause the mislocalisation of TDP-43 in human induced pluripotent stem cell derived motor neurons [44]. This indicates that not only can TDP-43 mislocalisation cause hyperexcitability, but this in turn may drive more TDP-43 into the cytoplasm in a positive feedback cycle.

The TDP-43^WT^ mouse used in these studies expresses TDP-43 with the same system as the TDP-43^ΔNLS^ mouse, yet TDP-43^WT^ layer V pyramidal neurons showed no changes to intrinsic or synaptic excitability compared to wild-type layer V pyramidal neurons. Neurodegeneration has previously been reported with overexpression of wild-type TDP-43 induced after weaning, and the rate of degradation becomes dependent on the expression levels of the promoter [45-47]. In the current study, the expression levels of TDP-43^WT^ did not amount to an overt physiological change, in contrast mislocalised TDP-43^ΔNLS^ expression on the same promoter system which caused broad functional alterations. This puts the subcellular localisation of TDP-43, rather than TDP-43 overexpression, at the forefront of TDP-43-mediated dysfunction in ALS.

## Conclusions

Our data addresses one of the long held questions regarding the pathogenesis of ALS; does TDP-43 mislocalisation, a key pathological feature of sporadic and familial ALS, cause hyperexcitability of neurons vulnerable to degeneration in ALS? In the present study, we observed functional changes in both the intrinsic excitability and the excitatory synaptic inputs to motor cortex layer V pyramidal neurons expressing wild-type, but mislocalised, TDP-43, as occurs in sporadic ALS. These data emphasise the importance of understanding neuronal excitability and synaptic changes as pathogenic events within the cascade of changes that are occurring in ALS. Therapeutics targeting cytoplasmic TDP-43 protein, along with therapies to ameliorate or modulate excitability should be strongly considered in the future.

## Declarations

### Ethics approval and consent to participate

All experiments were approved by the Animal Ethics Committee of the University of Tasmania (#A16593) in accordance with the Australian Code of Practice for the Care and Use of Animals for Scientific Purposes (2013).

### Consent for publication

Not applicable

### Availability of data and material

The current datasets are available from the corresponding author on reasonable request.

### Authors’ contributions

MD and CB wrote the manuscript. MD performed experiments. CB, AW, TD, AKW and KEL planned experiments. AW trained MD on electrophysiology. TD, AW, AKW, KEL edited the manuscript.

## Acknowledgements

University of Tasmania Animal facility staff for breeding and maintenance of mice used in these experiments.

## References

1. Al-Chalabi, A. and O. Hardiman, The epidemiology of ALS: a conspiracy of genes, environment and time. Nat Rev Neurol, 2013. 9(11): p. 617–28.

2. Neumann, M., Molecular neuropathology of TDP-43 proteinopathies. Int J Mol Sci, 2009.10(1): p. 232–46.

3. Neumann, M.S., DM. Kwong, LK. Truax, AC. Micsenyi, MC. Chou, TT. Bruce, J. Schuck, T. Grossman, M. Clark, CM. McCluskey, LF. Miller, BL. Masliah, E. Mackenzie, IR. Feldman, H. Feiden, W. Kretzschmar, HA. Trojanowski, JQ. Lee, VM., Ubiquitinated TDP-43 in frontotemporal lobar degeneration and amyotrophic lateral sclerosis. Science, 2006. 314: p. 130–133.

4. Ayala, Y.M., et al., TDP-43 regulates its mRNA levels through a negative feedback loop. EMBO J, 2011. 30(2): p. 277–88.

5. Liu-Yesucevitz, L., et al., Local RNA translation at the synapse and in disease. J Neurosci, 2011. 31(45): p. 16086–93.

6. Grieve, S.M., et al., Potential structural and functional biomarkers of upper motor neuron dysfunction in ALS. Amyotroph Lateral Scler Frontotemporal Degener, 2015. 17(1–2): p. 85–92.

7. Mills, K.R. and K.A. Nithi, Corticomotor threshold is reduced in early sporadic amyotrophic lateral sclerosis. Muscle Nerve, 1997. 20(9): p. 1137–1141.

8. Geevasinga, N., et al., Diagnostic utility of cortical excitability studies in amyotrophic lateral sclerosis. Eur J Neurol, 2014. 21(12): p. 1451–7.

9. Vucic, S., G.A. Nicholson, and M.C. Kiernan, Cortical hyperexcitability may precede the onset of familial amyotrophic lateral sclerosis. Brain, 2008. 131(Pt6): p. 1540–50.

10. Menon, P., M.C. Kiernan, and S. Vucic, Cortical hyperexcitability precedes lower motor neuron dysfunction in ALS. Clin Neurophysiol, 2015.126(4): p. 803–9.

11. Van den Bos, M.A.J., et al., Imbalance of cortical facilitatory and inhibitory circuits underlies hyperexcitability in ALS. Neurology, 2018.91(18): p. e1669–e1676.

12. Fogarty, M.J., P.G. Noakes, and M.C. Bellingham, Motor cortex layer V pyramidal neurons exhibit dendritic regression, spine loss, and increased synaptic excitation in the presymptomatic hSOD1(G93A) mouse model of amyotrophic lateral sclerosis. J Neurosci, 2015. 35(2): p. 643–7.

13. Saba, L., et al., Altered Functionality, Morphology, and Vesicular Glutamate Transporter Expression of Cortical Motor Neurons from a Presymptomatic Mouse Model of Amyotrophic Lateral Sclerosis. Cereb Cortex, 2016. 26(4): p. 1512–28.

14. Saba, L., et al., Modified age-dependent expression of NaV1.6 in an ALS model correlates with motor cortex excitability alterations. Neurobiol Dis, 2019. 130: p. 104532.

15. Handley, E.E., et al., Synapse Dysfunction of Layer V Pyramidal Neurons Precedes Neurodegeneration in a Mouse Model of TDP-43 Proteinopathies. Cereb Cortex, 2017. 27(7): p. 3630–3647.

16. Zhang, W., et al., Hyperactive somatostatin interneurons contribute to excitotoxicity in neurodegenerative disorders. Nat Neurosci, 2016.19(4): p. 557–559.

17. Igaz, L.M., et al., Dysregulation of the ALS-associated gene TDP-43 leads to neuronal death and degeneration in mice. J Clin Invest, 2011.121(2): p. 726–38.

18. Alfieri, J.A., N.S. Pino, and L.M. Igaz, Reversible behavioral phenotypes in a conditional mouse model of TDP-43 proteinopathies. J. Neurosci, 2014. 34(46): p. 15244–59.

19. Paxinos, G. and K. Franklin, The mouse brain in stereotaxic coordinates. Academic Press, 2007.

20. Ting, J.T., et al., Acute brain slice methods for adult and aging animals: application of targeted patch clamp analysis and optogenetics. Methods Mol Biol, 2014. 1183: p. 221–42.

21. Han, H.J., et al., Strain background influences neurotoxicity and behavioral abnormalities in mice expressing the tetracycline transactivator. J Neurosci, 2012. 32(31): p. 10574–86.

22. Harb, K., et al., Area-specific development of distinct projection neuron subclasses is regulated by postnatal epigenetic modifications. Elife, 2016. 5: p. e09531.

23. Arlotta, P., et al., Neuronal subtype-specific genes that control corticospinal motor neuron development in vivo. Neuron, 2005. 45(2):p. 207–21.

24. Rosier, K.T., A. Hess, CW. Magistris, MR., Quantification of upper motor neuron loss in amytrophic lateral sclerosis. Clin Neurophysiol, 2000. 111: p. 2208–2218.

25. Ravits, J.P., P. Jorg, C., Focality of upper and lower motor neuron degeneration at the clinical onset of ALS. Neurology, 2007. 68(19): p. 1571–1575.

26. Ravits, J. and A. LaSpada, ALS motor phenotype heterogeneity, focality, and spread. Neurology, 2009. 73: p. 805–811.

27. Rothstein, J.T., G. Kuncl, RW. Clawson, L. Cornblath, DR. Drachman, DB. Pestronk, A. Stauch, BL. Coyle, J., Abnormal excitatory amino acid metabolism in amyotrophic lateral sclerosis. Annals of Neurology, 1990. 28(1): p. 18–25.

28. Vucic, S., et al., Cortical excitability distinguishes ALS from mimic disorders. Clin Neurophysiol, 2011. 122(9): p. 1860–6.

29. Vucic, S. and M.C. Kiernan, Cortical excitability testing distinguishes Kennedy’s disease from amyotrophic lateral sclerosis. Clin Neurophysiol, 2008. 119(5): p. 1088–96.

30. Horoupian, D., et al., Dementia and motor neuron disease: morphometric, biochemical, and Golgi studies.. Ann. Neurol., 1984. 4(16): p. 305–316.

31. Zona, C., M. Pieri, and I. Carunchio, Voltage-dependent sodium channels in spinal cord motor neurons display rapid recovery from fast inactivation in a mouse model of amyotrophic lateral sclerosis. J Neurophysiol, 2006. 96(6): p. 3314–22.

32. Buskila, Y., et al., Dynamic interplay between H-current and M-current controls motoneuron hyperexcitability in amyotrophic lateral sclerosis. Cell Health and Disease, 2019. 10(4): p. 310–314.

33. Faber, E.S. and P. Sah, Functions of SK channels in central neurons. Clin Exp Pharmacol Physiol, 2007. 34(10): p. 1077–83.

34. Urbani, A. and O. Belluzzi, Riluzole inhibits the persistent sodium current in mammalian CNS neurons. European Journal of Neuroscience, 2000. 12: p. 3567–3574.

35. Geevasinga, N., et al., Riluzole exerts transient modulating effects on cortical and axonal hyperexcitability in ALS. Amyotroph Lateral Scler Frontotemporal Degener, 2016. 17(7–8): p. 580–588.

36. Kim, J., et al., Changes in the Excitability of Neocortical Neurons in a Mouse Model of Amyotrophic Lateral Sclerosis Are Not Specific to Corticospinal Neurons and Are Modulated by Advancing Disease. J Neurosci, 2017. 37(37): p. 9037–9053.

37. Anderson, C.T., et al., Sublayer-specific microcircuits of corticospinal and corticostriatal neurons in motor cortex. Nat Neurosci, 2010. 13(6): p. 739–44.

38. Fogarty, M.J., et al., Motor Areas Show Altered Dendritic Structure in an Amyotrophic Lateral Sclerosis Mouse Model. Front Neurosci, 2017. 11: p. 609.

39. Desai, N.S., N.J. Rutherford, and G.G. Turrigiano, Plasticity in the intrinsic excitability of cortical pyramidal neurons. Nature Neuroscience, 1999. 2(6): p. 515–520.

40. Turrigiano, G., Homeostatic synaptic plasticity: local and global mechanisms for stabilizing neuronal function. Cold Spring Harb Perspect Biol, 2012. 4(1): p. aOO5736.

41. Orr, B.O., et al., Presynaptic Homeostasis Opposes Disease Progression in Mouse Models of ALS-Like Degeneration: Evidence for Homeostatic Neuroprotection. Neuron, 2020.

42. Amlie-Wolf, A., et al., Transcriptomic Changes Due to Cytoplasmic TDP-43 Expression Reveal Dysregulation of Histone Transcripts and Nuclear Chromatin. PLoS One, 2015. 10(10): p. e0141836.

43. Sephton, C.F., et al., Identification of neuronal RNA targets of TDP-43-containing ribonucleoprotein complexes. J Biol Chem, 2011. 286(2): p. 1204–15.

44. Weskamp, K., et al., Shortened TDP-43 isoforms upregulated by neuronal hyperactivity drive TDP43 pathology in ALS. J Clin Invest, 2020. 130(3): p. 1139–1155.

45. Stallings, N.R., et al., Progressive motor weakness in transgenic mice expressing human TDP-43. Neurobiol Dis, 2010. 40(2): p. 404–14.

46. Xu, Y.F., et al., Wild-type human TDP-43 expression causes TDP-43 phosphorylation, mitochondrial aggregation, motor deficits, and early mortality in transgenic mice. J Neurosci, 2010. 30(32): p. 10851–9.

47. Swarup, V., et al., Pathological hallmarks of amyotrophic lateral sclerosis/frontotemporal lobar degeneration in transgenic mice produced with TDP43 genomic fragments. Brain, 2011. 134(Pt 9): p. 2610–26.

